# Differential EBV protein-specific antibody response between responders and non-responders to EBVSTs immunotherapy

**DOI:** 10.1101/2024.08.14.607997

**Authors:** Yomani D. Sarathkumara, Nathan W. Van Bibber, Zhiwei Liu, Helen E. Heslop, Rayne H Rouce, Anna E. Coghill, Cliona M Rooney, Carla Proietti, Denise L. Doolan

## Abstract

Epstein–Barr virus (EBV) is associated with a diverse range of lymphomas. EBV-specific T-cell (EBVST) immunotherapies have shown promise in safety and clinical effectiveness in treating EBV-associated lymphomas, but not all patients respond to treatment.

To identify the set of EBV-directed antibody responses associated with clinical response in patients with EBV-associated lymphomas, we comprehensively characterized the immune response to the complete EBV proteome using a custom protein microarray in 56 EBV-associated lymphoma patients who were treated with EBVST infusions enrolled in Phase I clinical trials.

Significant differences in antibody profiles between responders and non-responders emerged at 3 months post-EBVST infusion. Twenty-five IgG antibodies were present at significantly higher levels in non-responders compared to responders at 3 months post-EBVST infusion, and 10 of these IgG antibody associations remained after adjustment for sex, age, and cancer diagnosis type. Random forest prediction analysis further confirmed that these 10 antibodies were important for predicting clinical response. Differential IgG antibody responses were directed against LMP2A (four fragments), BGRF1/BDRF1 (two fragments), LMP1, BKRF2, BKRF4, and BALF5.

Paired analyses using blood samples collected at both pre-infusion and 3 months post-EBVST infusion indicated an increase in the mean antibody level for six other anti-EBV antibodies (IgG: BGLF2, LF1, BGLF3; IgA: BGLF3, BALF2, BBLF2/3) in non-responders. Overall, our results indicate that EBV-directed antibodies can be biomarkers for predicting the clinical response of individuals with EBV-associated lymphomas treated with EBVST infusions.

## Introduction

Epstein-Barr virus (EBV) is associated with a broad spectrum of hematologic malignancies, including Burkitt lymphoma, Hodgkin lymphoma, a subset of Non-Hodgkin lymphomas, including extranodal natural killer/T-cell lymphoma (NKTCL), and post-transplant lymphoproliferative disease (PTLD) (1). EBV-associated lymphomas are characterized by the presence of viral antigens that serve as potential targets for T-cell immunotherapy (2). Specifically, T-cells designed to target and respond to EBV-specific antigens (known as EBV-specific T-cells or EBVSTs) can exert immunostimulatory effects by secreting cytokines and chemokines upon target antigen recognition that recruit and activate local immune responses to viral and non-viral antigens associated with the tumor (3)(4).

EBV-associated malignancies exhibit specific expression patterns of a small set of EBV proteins that determine their EBV latency types (5, 6). Among these, Type III latency emerges only under conditions of severe immunosuppression and is more susceptible to adoptive T-cell therapy, likely because of the potent immunogenicity of Type III viral antigens (7–9). The first EBVSTs in the 1990s were donor-derived (LCL-activated) and administered in the hematopoietic stem cell transplant (HSCT) setting, an immunosuppressed patient setting. EBVSTs directed to viral antigens generated from off-the-shelf donors have effectively treated EBV-associated PTLD in the HSCT setting (10–12).

Type I/II latency tumors express EBV proteins, including EBV nuclear antigen (EBNA1) and latent membrane proteins (LMP 1 and 2), but they are less immunogenic compared to Type III tumors and thus potentially more challenging to treat with EBVSTs (2). However, T-cell immunotherapy against the LMP1 and/or LMP2 antigens was identified as a durable, safe, and effective approach without significant toxicity in Hodgkin lymphoma and NKTCL (i.e., Type II tumors) (4, 14, 15). Despite these promising clinical results, achieving long-term remission of type II latency lymphoma remains a challenge. Certain patient subgroups exhibit partial or no responses, often followed by relapses during T-cell immunotherapy (16). Previous research on EBVST immunotherapy under immunosuppressed conditions revealed instances of epitope spreading (2). This phenomenon, where the immune profile (i.e., viral antigens eliciting immune responses) extended beyond the initially targeted EBV antigens, was observed in patients who achieved positive clinical responses (4).

To better understand which EBV-directed humoral immune responses might be most indicative of clinical responses to EBVST infusions in EBV-associated lymphoma patients, the current study analyzed both IgG and IgA antibody responses against the complete EBV proteome. We employed a custom EBV protein microarray to measure both IgG and IgA antibodies against 202 EBV protein sequences in plasma samples from a cohort of 56 patients with EBV-associated lymphomas treated with autologous or allogeneic (third-party) EBVST infusions enrolled in Phase I clinical trials conducted by the Baylor College of Medicine, USA.

Unlike gene expression analyses in tumors, which may require invasive procedures such as biopsies, measuring antibodies can serve as non-invasive, informative biomarkers assessing patient-level EBV response to gain insights into the host immune response. Hence, antibodies could serve as indicators for patient monitoring and clinical management of EBV-associated lymphomas. In addition, these antibody-directed EBV antigens could guide development of immunotherapy targets for treating EBV-associated lymphomas.

## Materials and Methods

### Subjects and samples

Archived plasma samples were selected from patients diagnosed with EBV-positive lymphomas receiving infusions of autologous or allogeneic (third-party) peptide-stimulated EBV-specific T-cells (EBVSTs) collected from one of three multi-centre Phase I clinical trials: GRALE (NCT01555892 in which autologous EBVSTs were infused), PREVALE (NCT02973113 in which autologous EBVSTs were infused with Nivolumab) and MABEL (NCT02287311 in which banked allogeneic EBVSTs were infused), all conducted at the Baylor College of Medicine (Table 1). These studies were approved by the Institutional Review Boards of the Baylor College of Medicine, the National Cancer Institute (NCI), representative institutes, and affiliated hospitals and conducted under an Investigational New Drug application to the FDA. Written informed consent was obtained from all participants. All laboratory testing of archived samples was conducted under a protocol approved by the James Cook University Human Research Ethics Committee (H7696).

**Table 1.** Baseline characteristics of all individuals and by response status (responders versus non-responders) to EBVSTs treatment.

| Characteristic | Overall,<br>N = 56 (%) | Treatment Response |  | *p*-value<sup>†</sup> |
| --- | --- | --- | --- | --- |
|  |  | Responders,<br>N = 36 (64%) | Non-responders,<br>N = 20 (36%) (%) |  |
| Sex |  |  |  |  |
| Female | 17 (30%) | 10 (28%) | 7 (35%) | 0.6 |
| Male | 39 (70%) | 26 (72%) | 13 (65%) |  |
| Age Groups |  |  |  |  |
| 0-30 | 24 (43%) | 17 (47%) | 7 (35%) | 0.6 |
| 31-60 | 21 (38%) | 12 (33%) | 9 (45%) |  |
| 61-100 | 11 (20%) | 7 (19%) | 4 (20%) |  |
| Ethnic Groups |  |  |  |  |
| Hispanic | 12 (21%) | 9 (25%) | 3 (15%) | 0.5 |
| Non-Hispanic | 44 (79%) | 27 (75%) | 17 (85%) |  |
| Diagnosis |  |  |  |  |
| T-cell lymphomas | 13 (23%) | 8 (22%) | 5 (25%) | 0.7 |
| Hodgkin disease | 19 (34%) | 14 (39%) | 5 (25%) |  |
| Burkitt/Large Cell |  |  |  |  |
| Lymphoma | 13 (23%) | 7 (19%) | 6 (30%) |  |
| Other | 11 (20%) | 7 (19%) | 4 (20%) |  |

<sup>†</sup>Pearson's Chi-squared test; Fisher's exact test.

The 56 patients selected for our study were all diagnosed with EBV-positive lymphoma, were treated with EBVSTs per clinical trial protocols, and had residual, banked blood samples available for correlate biomarker testing. They were categorized as positive responders (complete or partial response recorded at last follow-up; n=36) or non-responders (progressive or stable disease recorded at last follow-up; n=20). The samples were collected at the pre-EBVST infusion time point and three post-infusion time points (2-weeks, 4-weeks, and 3-months).

### EBV custom protein microarray

Plasma samples were probed using a custom EBV protein microarray targeting IgA and IgG antibodies against 202 EBV protein sequences representing the complete EBV proteome, as previously described (17–21). Briefly, our comprehensive EBV protein microarray contains 199 EBV protein sequences generated from five different EBV strains (AG876, Akata, B95-8, Mutu, and Raji) representing nonredundant open reading frames and predicted splice variants from 86 EBV proteins. Also included on the array were peptide sequences representing three synthetic EBV peptides (VCAp18, EBNA-1, and EA p47) considered putative cancer biomarkers (17). Four “noDNA” (non-translated protein) spots were included to correct for person-specific background.

Plasma samples from each study participant (**Table 1**) were tested, blinded on response status, as described previously (17, 19–21). Briefly, antibody responses were detected with biotin-conjugated goat anti-human IgG (1:1000 dilution) or IgA (1:500 dilution) secondary antibodies (Jackson ImmunoResearch Laboratories, West Grove, PA, USA) and visualized with streptavidin-conjugated SureLight^®^ P3 (Columbia Biosciences, Columbia, MD, USA) (1:200 dilution) antibody. After probing, air-dried probed slides were scanned on an Axon GenePix 4300A (Molecular Devices). Raw fluorescence intensities were corrected for spot-specific background using the Axon GenePix Pro 7 software, and data were variant log-transformed using variance stabilizing normalization (VSN) transformation in Gmine (22). The array output was then standardized, referred to as the standardized signal intensity (SSI), to the person-specific background using the individual’s cut-off (mean ±1.5 SD of the four “no DNA” spots). Positivity was defined as a standardized signal intensity > 1.0, and output was further categorized into positive (1) and negative (0) responses.

Nineteen duplicate samples were included for assessment of reproducibility, and a cut-off coefficient of variation [CV] of 30% was selected (19–21). We excluded array spots with CVs ≤ 30% from the analysis, leaving 74 IgG and 202 IgA markers for comparisons between responders and non-responders.

### Statistical analysis

All statistical analyses were performed using R Statistical Software (v4.2.1; R Core Team 2022)(23). In addition to reporting nominal *p*-values from primary statistical tests, the Benjamini and Hochberg false discovery rate (FDR=5%) method was applied to account for multiple testing.

Differences in the distribution of demographic and clinical variables (sex, age group, ethnic group, diagnosis) between responders and non-responders were assessed using a chi-square test. Differences in the mean standardized signal intensity (SSI) for IgG and IgA antibodies between the responders (n=36) and non-responders (n=20) were assessed using unpaired t-tests at each blood collection timepoint. The odds ratios (ORs) and 95% CIs for the association between detection of each anti-EBV antibody (*i.e*., categorized into binary responses: positive=1, negative=0) and treatment response status (positive response /responders=1, no response, non-responders=0) were calculated using logistic regression models adjusted for sex, age at enrolment, and diagnosis type (T-cell lymphomas, Hodgkin’s lymphoma, B-cell lymphomas, and other lymphoma). In addition to the adjusted logistic regression models, the continuous SSI output at the 3-month post-infusion timepoint was analyzed with random forest models (24) using the R package random forest (v4.7.1) (25) to rank IgG or IgA antibodies based on their importance in differentiating samples into the defined groups (responders/non-responders). Specifically, antibodies at each time point were ranked using the Mean Decrease in Gini index (MDG) and Mean Decrease in Accuracy (MDA) metrics, which order variables according to their importance in improving prediction model purity and accuracy, respectively.

The total change in antibody (Ab) response between the pre-infusion and 3-month post-infusion time points (Ab_3mo_−Ab_pre_) was evaluated among responders and non-responders for patients with matched blood samples at both time points (n = 41 samples for IgG [30 responders, 11 non-responders]; n = 42 samples for IgA [30 responders, 12 non-responders]). The odds ratios (ORs) and 95% CIs for the association between each anti-EBV antibody variable (*i.e*., Ab_3mo_−Ab_pre_ change in SSI) and treatment response status were calculated using logistic regression models adjusted for sex, age at enrolment, and diagnosis type (T-cell lymphomas, Hodgkin’s lymphoma, B-cell lymphomas, and other lymphomas).

## Results

### Population characteristics

All patients included in the EBVST immunotherapy study were diagnosed with EBV-associated lymphoma (n=56) and classified as either responders (n=36) or non-responders (n=20) at the end of clinical trial follow-up (**Table 1**). No differences in sex ( *p*= 0.6), age (*p*=0.6), race/ethnicity(*p*=0.8) or lymphoma diagnosis category (*p*=0.7) were observed between responders and non-responders.

### EBV proteome-wide analysis of virus-directed humoral response pre- and post-EBVSTinfusions

We evaluated differences in the mean standardized signal intensity (SSI) for 74 IgG and 202 IgA antibodies between responders (n=36) and non-responders (n=20) to EBVST immunotherapy by unpaired-t-tests at pre-clinical treatment (baseline) and at three time points after treatment (2-weeks, 4-weeks, and 3-month) (**Figure 1**, **Table 2**). Pre-treatment, IgG antibodies against EBV nuclear antigen leader protein (EBNA-LP) (nominal *p* =0.03, t-test) and IgA antibodies for lytic gene BGLF3.5 (nominal *p* =0.04, t-test) were higher among the non-responders than the responders to EBVST immunotherapy (**Table 2**).

**Figure 1.**
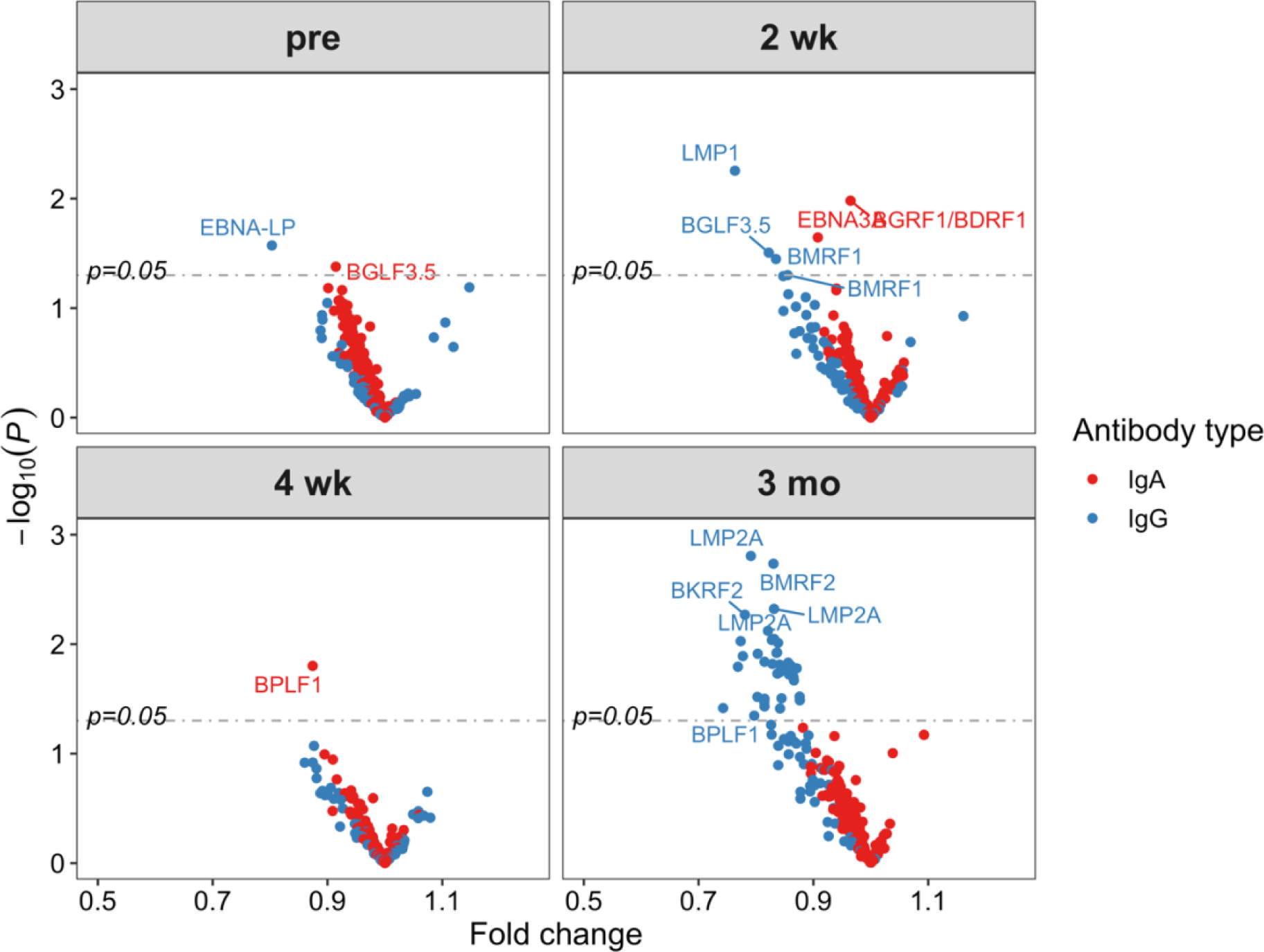
Differential IgA and IgG reactivity between responders and non-responders to EBVST immunotherapy. Differences in the mean antibody response in responders *vs*. non-responders to EBVST immunotherapy at pre-infusion and 2-weeks (2 wk), 4-weeks (4 wk), and 3-months (3 mo) post-infusion. The x-axis of the volcano plot displays the fold change (ratio of standardized signal intensity [SSI] comparing responders *vs*. non-responders) for all antibodies with CV < 30% (IgG=74, IgG= 202) (red, IgA; blue, IgG). The y-axis illustrates the *p*-value corresponding to the t-test for a difference in SSI between treatment responders and non-responders. The dashed lines represent the nominally significant *p*-value threshold.

**Table 2.** Table of EBV proteins on microarrays (microarray sequence, protein name, and EBV life cycle) for IgG and IgA antibodies with evidence of differential levelsbetween responders and non-responders to EBVST immunotherapy, by time point post-infusion.

| Array sequence | Protein name | Life cycle | T-test *p*-value | FDR adj *P*-value | Responder group Mean | Non-responder group Mean | IgG/ IgA |
| --- | --- | --- | --- | --- | --- | --- | --- |
| **Pre-EBVST immunotherapy** |  |  |  |  |  |  |  |
| YP_001129440.1-20824-20955 | EBNA-LP | Latent | 0.03 | 0.998 | 1.04 | 1.29 | IgG |
| YP_001129483.1-112496-112035 | BGLF3.5 | Late lytic | 0.04 | 0.903 | 1.15 | 1.26 | IgA |
| **2-weeks post-EBVST immunotherapy** |  |  |  |  |  |  |  |
| YP_001129485.1-117754-118890 | BGRF1/BDRF1 | Late lytic | 0.01 | 0.998 | 0.97 | 1.01 | IgA |
| AFY97906.1-168167-168081 | LMP1 | Latent | 0.01 | 0.411 | 0.99 | 1.30 | IgG |
| YP_001129463.1-80447-82888 | EBNA3A | Latent | 0.02 | 0.998 | 1.23 | 1.36 | IgA |
| YP_001129483.1-112496-112035 | BGLF3.5 | Late lytic | 0.03 | 0.719 | 0.76 | 0.92 | IgG |
| AFY97929.1-67486-68700 | BMRF1 | Early lytic | 0.04 | 0.719 | 1.83 | 2.19 | IgG |
| YP_001129454.1-67745-68959 | BMRF1 | Early lytic | 0.05 | 0.719 | 1.97 | 2.30 | IgG |
| **4-weeks post-EBVST immunotherapy** |  |  |  |  |  |  |  |
| YP_001129449.1-59370-49906-3 | BPLF1 | Late lytic | 0.02 | 0.999 | 0.86 | 0.99 | IgA |
| **3-months post-EBVST immunotherapy** |  |  |  |  |  |  |  |
| YP_001129436.1-871-951 | LMP2A | Latent | 0.00 | 0.058 | 1.10 | 1.39 | IgG |
| YP_001129455.1-68964-70037 | BMRF2 | Glycoprotein | 0.00 | 0.058 | 1.14 | 1.38 | IgG |
| YP_001129436.1-360-458 | LMP2A | Latent | 0.00 | 0.058 | 1.09 | 1.31 | IgG |
| YP_001129472.1-98500-98913 | BKRF2 | Glycoprotein | 0.01 | 0.058 | 0.92 | 1.18 | IgG |
| YP_001129436.1-540-788 | LMP2A | Latent | 0.01 | 0.058 | 0.94 | 1.15 | IgG |
| YP_001129484.1-113481-112483 | BGLF3 | Late lytic | 0.01 | 0.058 | 1.31 | 1.58 | IgG |
| YP_001129506.1-154125-153187 | BILF1 | Glycoprotein | 0.01 | 0.058 | 1.14 | 1.37 | IgG |
| YP_001129479.1-107679-108896 | BBRF3 | Glycoprotein | 0.01 | 0.058 | 1.53 | 1.97 | IgG |
| YP_401715.1-160908-158851 | BALF3 | Late lytic | 0.01 | 0.058 | 1.23 | 1.46 | IgG |
| CAA24827.1-122341-120929 | BGLF5 | Early lytic | 0.01 | 0.058 | 0.91 | 1.09 | IgG |
| YP_001129461.1-76771-77259 | BLRF2 | Late lytic | 0.01 | 0.058 | 2.18 | 2.61 | IgG |
| CAA24860.1-102116-101445 | BZLF2 | Glycoprotein | 0.01 | 0.058 | 1.95 | 2.43 | IgG |
| YP_001129485.1-117754-118890 | BGRF1/BDRF1 | Late lytic | 0.01 | 0.058 | 0.87 | 1.11 | IgG |
| YP_001129466.1-90630-89959 | BZLF2 | Glycoprotein | 0.01 | 0.058 | 2.02 | 2.47 | IgG |
| YP_001129500.1-136454-135636 | BVLF1 | Late lytic | 0.01 | 0.058 | 1.24 | 1.45 | IgG |
| YP_001129504.1-151808-150519 | LF2 | Early lytic | 0.02 | 0.058 | 1.14 | 1.38 | IgG |
| YP_001129436.1-1026-1196 | LMP2A | Latent | 0.02 | 0.058 | 1.05 | 1.24 | IgG |
| AFY97988.1-166888-166706 | BNLF2A | Late lytic | 0.02 | 0.058 | 1.17 | 1.36 | IgG |
| YP_001129440.1-35441-35473 | EBNA-LP | Latent | 0.02 | 0.058 | 0.83 | 1.09 | IgG |
| YP_001129470.1-94844-96457 | BRRF2 | Late lytic | 0.02 | 0.058 | 2.15 | 2.54 | IgG |
| YP_001129496.1-131574-129454 | BXLF2 | Glycoprotein | 0.02 | 0.058 | 1.35 | 1.54 | IgG |
| YP_001129498.1-133398-134144 | BXRF1 | Late lytic | 0.02 | 0.058 | 1.12 | 1.33 | IgG |
| YP_001129486.1-115415-114405 | BGLF2 | Early lytic | 0.02 | 0.058 | 1.16 | 1.39 | IgG |
| YP_001129439.1-9659-10171 | BcRF1 | Late lytic | 0.02 | 0.058 | 1.12 | 1.31 | IgG |
| YP_001129512.1-166530-167195 | BARF1 | Early lytic | 0.02 | 0.059 | 1.13 | 1.31 | IgG |
| CAA24838.1-61507-62037 | BFRF3 | Late lytic | 0.02 | 0.061 | 2.25 | 2.59 | IgG |
| YP_001129448.1-49335-49865 | BFRF3 | Late lytic | 0.03 | 0.078 | 2.37 | 2.70 | IgG |
| CAA24829.1-124938-125915 | BGRF1/BDRF1 | Late lytic | 0.03 | 0.078 | 0.92 | 1.15 | IgG |
| YP_001129488.1-117560-116883 | BDLF4 | Early lytic | 0.03 | 0.078 | 1.48 | 1.75 | IgG |
| YP_001129438.1-1736-5692-2 | FGAM | Other/Unknown | 0.03 | 0.078 | 1.35 | 1.65 | IgG |
| AFY97924.1-49199-49729 | BFRF3 | Late lytic | 0.03 | 0.078 | 2.27 | 2.59 | IgG |
| YP_001129489.1-117772-117539 | BDLF3.5 | Glycoprotein | 0.03 | 0.078 | 0.91 | 1.11 | IgG |
| AFY97906.1-168167-168081 | LMP1 | Latent | 0.04 | 0.082 | 1.09 | 1.34 | IgG |
| YP_001129440.1-20824-20955 | EBNA-LP | Latent | 0.04 | 0.082 | 0.97 | 1.31 | IgG |
| YP_001129454.1-67745-68959 | BMRF1 | Early lytic | 0.04 | 0.082 | 1.98 | 2.36 | IgG |
| CAA24839.1-71527-62078-3 | BPLF1 | Late lytic | 0.05 | 0.093 | 0.76 | 0.96 | IgG |

FDR correction method: Benjamini and Hochberg. The table is ordered by t-test *p*-value (lowest to highest). T-test *p*-values <0.05 is considered nomially significant for this analysis, with FDR-corrected values considered statistically significant.

At 2-weeks after immunotherapy treatment, IgG antibodies against LMP1 (nominal *p* =0.01, t-test), BGLF3.5 (nominal *p* =0.03, t-test), and two variants of BMRF1 (nominal *p* =0.04, and *p* =0.05, t-test), as well as IgA antibodies against BGRF1/BDRF1 (nominal *p* =0.01, t-test) and EBNA3A (nominal *p* =0.02, t-test), were elevated in non-responders. At 4-weeks post-treatment, IgA antibodies against BPLF1were higher in non-responders (nominal *p* = 0.02, t-test).

More pronounced IgG responses, but not IgA, were evident at 3-months post-EBVST infusion in non-responders (**Figure 1**). A total of 36 IgG antibodies were higher among non-responders (nominal *p* <0.05, t-test) (**Table 2**). When adjusted for multiple testing using the Benjamini and Hochberg false discovery rate (FDR), 25 IgG antibodies at the 3-month post-infusion timepoint reached borderline significance (FDR=0.058). No other marker/timepoint combinations reached significance after FDR correction.

Logistic regression models adjusted for sex, age, and diagnosis type were then applied to the antibodies with a nominal *p*-value < 0.05. (**Table S1)**. At 3-months post-treatment, a total of 10 IgG antibodies (eight of the 36 IgG antibodies identified by t-test and two additional IgG antibodies identified by logistic regression) were present at higher levels in non-responders (nominal *p* <0.05, logistic regression) (**Table S1)**.At 3 months post-EBVST infusion, Random Forest analysis identified informative predictors of the outcome of clinical response IgG antibodies using the Mean Decrease in Gini (MDG) and Mean Decrease in Accuracy (MDA) metrics (25). Antibodies scoring high for both metrics, indicating their status as important, were visualized on a multi-way importance plot (**Figure 2**). The ten antibodies significant at the 3-month time-point (by nominal *p*-value) in the logit models were also identified as important variables in the random forest model, demostrating robust agreement between the two methods. These IgG antibody responses were directed against LMP2A (4 fragmenst), BGRF1/BDRF1 (2 fragments), LMP1, BKRF2, BKRF4 and BALF5.

**Figure 2.**
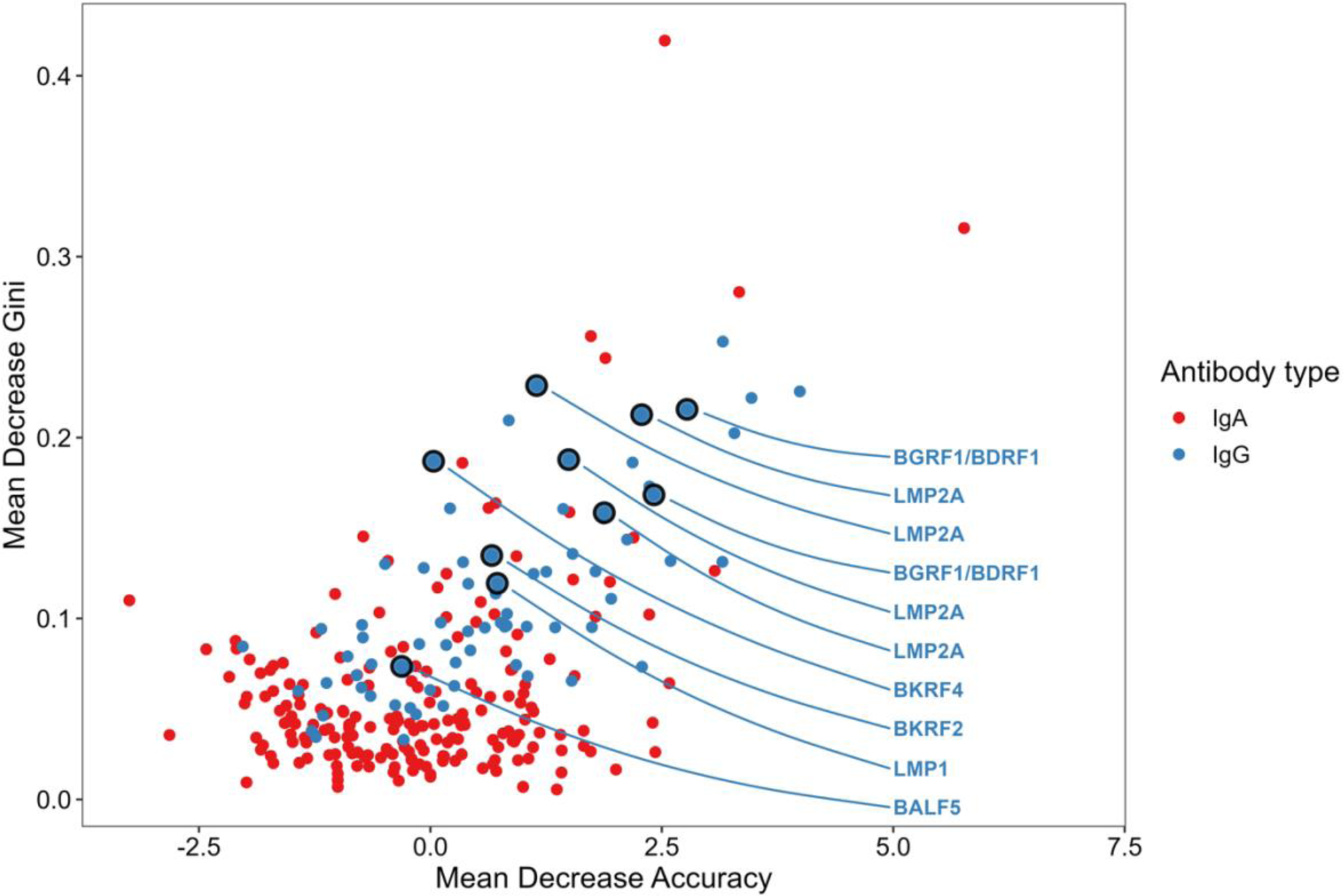
multi-way variable importance plot at 3 months post-infusion (mean decrease accuracy and mean decrease Gini). The higher the value of Mean Decrease Accuracy or Mean Decrease Gini score, the higher the importance of the variable in the model. Therefore, antibodies in the upper-right of the plot indicate those determined as important by both metrics. All the antibodies that were significant at the 3-month time-point (by nominal *p*-value) in the logit models are identified in the random forest as the most important variables, demostrating robustagreement between the two.

### Changes in EBV-directed antibody response during EBVST infusion treatment

Total change in antibody response between pre-infusion and 3-months post-infusion (Ab_3mo_−Ab_pre_) was evaluated among responders and non-responders. Only individuals with paired IgG (n=41,) and paired IgA data (n=42) at both time points were included in the analysis.

Total change in six anti-EBV antibodies (3 IgA; BGLF3, BALF2, BBLF2/3 and 3 IgG; BGLF2, LF1, BGLF3) was associated with treatment response (nominal *p* <0.05, logistic regression) when adjusted for sex, age, and diagnosis (**Table 3**); however, none of these retained significance after FDR correction. Ribbon plots in **Figure 3** show the changes in average responses per each antibody marker. Notably, the non-responder group exhibited increased mean antibody responses at 3-months compared to pre-treatment for all six antibody markers, indicating the elicitation of robust antibody responses for EBVST infusions in individuals who did not clinically respond to treatment (non-responders).

**Figure 3.**
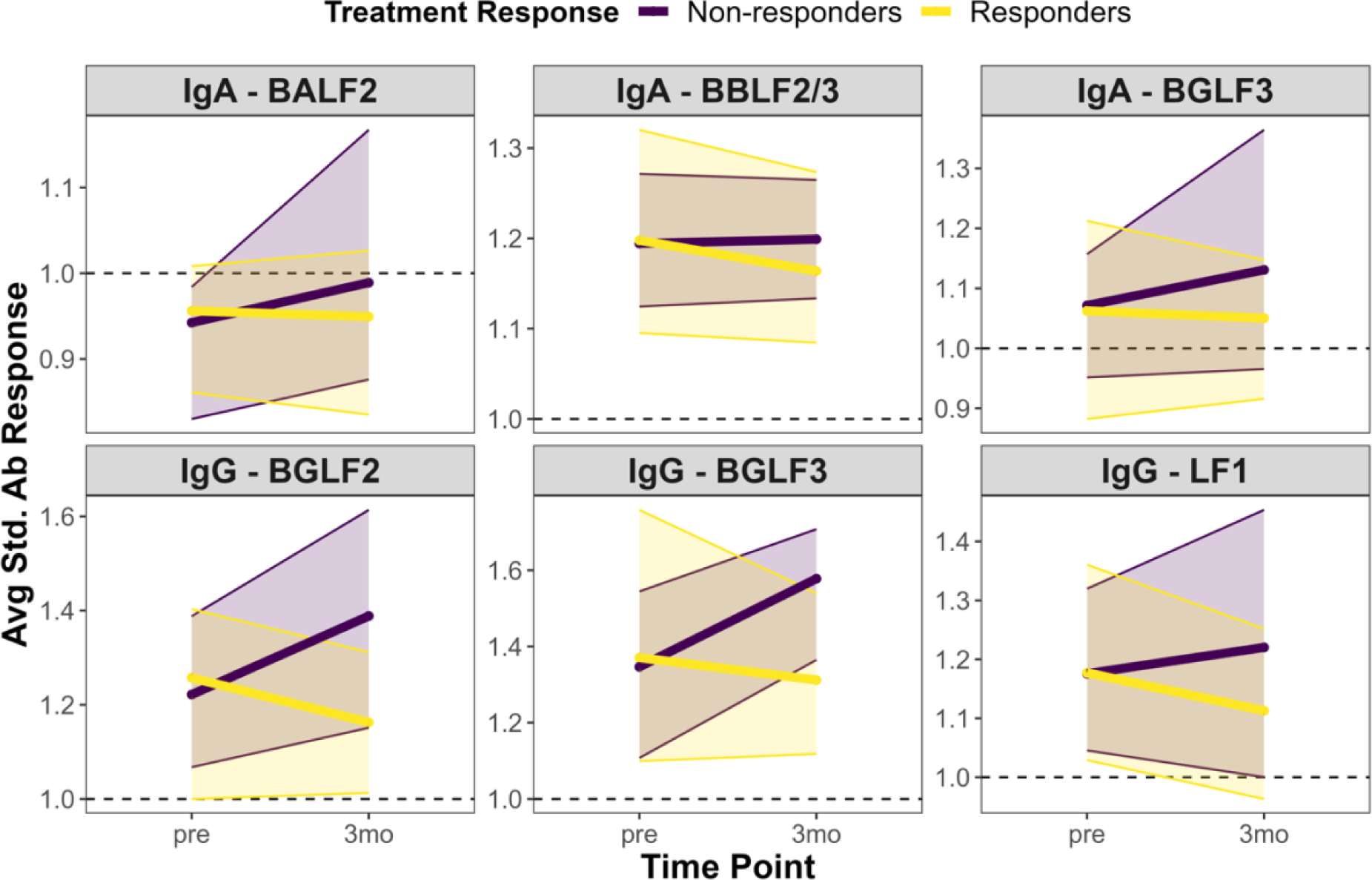
Change in antibody response between pre-infusion and 3-months post-infusion. The ribbon plots show the 3-month change in the mean (and 25^th^ to 75^th^ percentile range) antibody response (SSI) for EBV antibody markers with nominal *p* <0.05 in adjusted logistic regression models. Purple indicates the non-responder group (N=20), while yellow indicates the responder group (N=36).

**Table 3:** Table of odds ratios for EBV-directed antibodies with evidence of an association between clinical response and the antibody change between the pre-infusion and 3-month post-infusion (Ab_3mo_−Ab_pre_) timepoints.

| Array sequence | Protein name | Life cycle | IgG/<br>IgA | Log. reg *p*-value | *p*-value (FDR) | OR (95% CI) |
| --- | --- | --- | --- | --- | --- | --- |
| YP_001129484.1-113481-112483 | BGLF3 | Late lytic | IgA | 0.036 | 0.889 | 0.011 (0-0.564) |
| YP_001129510.1-165796-162410-2 | BALF2 | Early lytic | IgA | 0.036 | 0.889 | 0 (0-0.154) |
| AFY97950.1-104968-104363 | BBLF2/3 | Early lytic | IgA | 0.043 | 0.889 | 0 (0-0.324) |
| YP_001129486.1-115415-114405 | BGLF2 | Early lytic | IgG | 0.042 | 0.359 | 0.06 (0.002-0.681) |
| YP_001129505.1-153178-151769 | LF1 | Other/Unknown | IgG | 0.045 | 0.359 | 0.036 (0.001-0.7) |
| YP_001129484.1-113481-112483 | BGLF3 | Late lytic | IgG | 0.046 | 0.359 | 0.075 (0.004-0.713) |

FDR correction method: Benjamini and Hochberg. The table is ordered by *p*-value (lowest to highest). Logistic regression *p*-values <0.05 were considered nomially significant for this analysis.

One unique observation was the consistent association between total change in both IgG (*p*=0.046, OR=0.075, 95% CI = 0.004, 0.713) and IgA (*p*=0.036, OR=0.011, 95% CI = 0, 0.564) antibody responses to BGLF3 and clinical non-response. BGLF3-IgG was also elevated at the 3-month timepoint in the non-responder group in univariate analyses (nominal *p* <0.05, t-test) and was identified as an important marker by random forest metrics (**Table S1**).

## Discussion

We observed notable differences in EBV-directed antibody responses at the 3-month post-EBVST infusion time point between responders and non-responders to EBVST immunotherapy. Specifically, IgG antibody responses against LMP2A (four fragments), BGRF1/BDRF1 (two fragments), LMP1, BKRF2, BKRF4, and BALF5 remained significantly associated with non-responseafter adjustment for sex, age, and cancer diagnosis and were also classified as important for distinguishing responders and non-responders for EBVST infusions using random forest prediction metrics.

EBV-positive tumors express viral latency-associated antigens that can be targeted for T-cell immunotherapy. The adoptive transfer of EBVSTs has proven to be a promising treatment option for immunogenic type III latency-derived PTLD, commonly occurring in transplant recipients with compromised immune systems (12). While previous studies have explored peripheral autoantibodies against tumor-associated proteins as markers for cancer and predictors of clinical outcomes (30–32), recent research on EBVSTs has primarily focused on a limited set of viral antigens expressed during latency, such as LMP1, LMP2, and EBNA1 (28).

Broadening the spectrum of viral antigens by proteome-wide profiling can be a promising approach for identifying which EBV-directed antigens might best predict clinical responses for EBV-associated lymphoma and for identifying an expanded set of target antigens for EBVSTs to help improve complete response rates for EBV-associated lymphomas treated using this type of immunotherapy. To address this, we utilized a custom protein microarray consisting of predicted sequences from the complete EBV proteome to measure IgG and IgA antibody responses. To our knowledge, this is the first report comprehensively evaluating patterns of anti-EBV antibody responses in EBV-positve lymphoma patients undergoing EBVST infusions.

Overall, our findings reveal that patients with positive clinical outcomes to EBVST immunotherapy were characterized by a low level of anti-EBV antibody responses, whereas patients who did not respond to treatment had notably higher anti-EBV antibodies, particularly at 3 months post-treatment. Antibodies serve as indicators of exposure, and elevated antibody responses in individuals who did not respond to EBVST infusions may suggest that they are experiencing uncontrolled EBV infection. Although we did not directly measure EBV levels in circulation, antibodies are metrics of the immune response to systemic exposure. Our data are consistent with previous reports showing persistently high EBV viremia levels in PTLD patients who poorly responded to immunotherapy (44, 45) but offer a more nuanced assessment of a specific set of EBV antigens whose lack of control is associated with poor clinical outcome. IgG antibody targeting multiple LMP2A sequences differed by clnical response.

LMP2A is a facilitator of B-cell survival, promoting viral persistence and supporting B-cell activation and transformation (30–33). LMP1 expression is identified as a surrogate marker of EBV in Hodgkin lymphoma and, as an oncogene, contributes to cell proliferation and survival in EBV-associated malignancies while suppressing apoptosis (29–31). As such, elevated expression in non-responders is biologically plausible and warrants further, refined targeting efforts in EBVST therapeutic approaches.

Multiple associations between elevated antibody response and clinical non-response were also observed for BGRF1/BDRF1, a gene set encoding DNA packaging terminase subunit that is believed to play a significant role in capsid assembly and/or stability and in nuclear egress (32, 33).

Also noteworthy antibodies are the three IgG (BGLF2, LF1, BGLF3) and three IgA (BGLF3, BALF2, BBLF2/3) identified as nominally elevated between the pre-infusion and 3 months post-infusion timepoints, as well as being associated with non-response. BGLF3 was of particular note given its consistent association with clinical response, not just in relation to antibody change but also in relation to the total level measured 3 months post-EBVST infusion; BGLF3 is part of a viral pre-initiation complex (vPIC) essential for regulating the expression of late EBV gene expression (37).

Beyond the fact that antibodies reflect exposure to systemic EBV infection, the contribution of B-cells and B-cell-mediated antibody responses, representing the humoral arm of the adaptive immune system, to anti-tumor activity and immunotherapies is not extensively explored in immune-oncology research (46). While B-cells are known to play roles in tumor immunology, including antigen presentation and producing tumor-specific antibodies (47, 48), their functions in facilitating anti-tumor responses, including antibody-dependent cell cytotoxicity (ADCC) and complement cascade activation, are not fully elucidated (49). (56, 57). Th2-mediated immunity has been traditionally considered to facilitate tumor growth by promoting angiogenesis and by inhibiting cell-mediated immunity, with several studies supporting that humoral immunity can have a negative impact on anti-tumor immunity (58, 59). Therefore, it is possible that EBVSTs promoted Th2 responses rather than Th1 responses in the non-responsive individuals.

One limitation of our study is the small sample size, which was dictated by the phase 1 clinical study designs. Consequently, it is crucial to validate and extend our initial observations in larger patient cohorts derived from clinical immunotherapy trials. In addition, we acknowledge the existence of several compartments of EBV-infected cells in the body, including EBV-infected B-cells in lymphoid tissues, EBV-infected epithelial cells in the oropharynx, and EBV-infected tumor cells. The antibody response that we measured in blood likely reflects a combination of these but is not a direct measurement of any given compartment. Whereas blood-based metrics can serve as accessible biomarkers, we note that further, tumor-based studies are needed to understand if any of our observations reflects the viral proteins expressed in the tumor.

In summary, this study provides the first attempt to examine the antibody responses against the complete EBV proteome in patients with EBV-associated lymphomas treated with EBVST immunotherapy. Our findings revealed a consistently elevated antibody response to EBV antigens in those with non-response to EBVST immunotherapy. We propose that the identified set of elevated anti-EBV antibodies are predictive markers of antigen persistence and clinical outcome post-EBVST infusion in those diagnosed with EBV-associated lymphomas. Further research, particularly in larger patient cohorts, is imperative to validate and extend these initial observations, enhancing our understanding of the underlying mechanisms and optimizing therapeutic strategies.

## Acknowledgments

This study was supported by the Intramural Research Program of the National Cancer Institute (NCI), and by a SPORE in Lymphooma P50CA126752 (HEH, CMR). DLD was supported by the National Health and Medical Research Council of Australia (NHMRC) Principal Research Fellowship (#1137285). YDS was supported by a Postgraduate Research Scholarship from James Cook University. We are grateful to the study participants without whom this work would not be possible.

## Author Contribution

YDS performed the protein microarray studies. YDS and NVB performed statistical analyses. YDS, NVB, AEC, DLD and CP wrote the manuscript. CMR and HEH designed the clinical research study. CMR, HEH and AEC selected the samples for laboratory study. DLD and AEC designed and supervised this study and acquired funding. DLD, CP and AES designed and supervised statistical analyses. YDS, NVB, AEC, CP and DLD interpreted data and critically reviewed the manuscript. All authors have reviewed and approved the final manuscript.

## References

1. Hudnall SD, Ge Y, Wei L, Yang N-P, Wang H-Q, Chen T. Distribution and phenotype of Epstein–Barr virus-infected cells in human pharyngeal tonsils. Modern Pathology. 2005;18(4):519–27.

2. Heslop HE, Sharma S, Rooney CM. Adoptive T-Cell Therapy for Epstein-Barr Virus-Related Lymphomas. Journal of Clinical Oncology. 2021;39(5):514–24.

3. Sharma S, Leung WK, Heslop HE. Virus-specific T cells for malignancies - then, now and where to? Current Stem Cell Reports. 2020;6(2):17–29.

4. Bollard CM, Gottschalk S, Torrano V, Diouf O, Ku S, Hazrat Y, et al. Sustained complete responses in patients with lymphoma receiving autologous cytotoxic T lymphocytes targeting Epstein-Barr virus latent membrane proteins. Journal of Clinical Oncology. 2014;32(8):798–808.

5. Middeldorp JM, Brink AATP, van den Brule AJC, Meijer CJLM. Pathogenic roles for Epstein–Barr virus (EBV) gene products in EBV-associated proliferative disorders. Critical Reviews in Oncology/Hematology. 2003;45(1):1–36.

6. Bauer M, Jasinski-Bergner S, Mandelboim O, Wickenhauser C, Seliger B. Epstein-Barr Virus-Associated Malignancies and Immune Escape: The Role of the Tumor Microenvironment and Tumor Cell Evasion Strategies. Cancers (Basel). 2021;13(20).

7. Gottschalk S, Rooney CM. Adoptive T-Cell Immunotherapy. Current Topics in Microbiology and Immunology. 2015;391:427–54.

8. Rooney CM, Ng CYC, Loftin S, Smith CA, Li C, Krance RA, et al. Use of gene-modified virus-specific T lymphocytes to control Epstein-Barr-virus-related lymphoproliferation. The Lancet. 1995;345(8941):9–13.

9. Rooney CM, Smith CA, Ng CYC, Loftin SK, Sixbey JW, Gan Y, et al. Infusion of Cytotoxic T Cells for the Prevention and Treatment of Epstein-Barr Virus–Induced Lymphoma in Allogeneic Transplant Recipients. Blood. 1998;92(5):1549–55.

10. Heslop HE, Slobod KS, Pule MA, Hale GA, Rousseau A, Smith CA, et al. Long-term outcome of EBV-specific T-cell infusions to prevent or treat EBV-related lymphoproliferative disease in transplant recipients. Blood. 2010;115(5):925–35.

11. Doubrovina E, Oflaz-Sozmen B, Prockop SE, Kernan NA, Abramson S, Teruya-Feldstein J, et al. Adoptive immunotherapy with unselected or EBV-specific T cells for biopsy-proven EBV+ lymphomas after allogeneic hematopoietic cell transplantation. Blood, The Journal of the American Society of Hematology. 2012;119(11):2644–56.

12. Bollard CM, Rooney CM, Heslop HE. T-cell therapy in the treatment of post-transplant lymphoproliferative disease. Nature Reviews Clinical Oncology. 2012;9(9):510–9.

13. Haque T, Wilkie GM, Taylor C, Amlot PL, Murad P, Iley A, et al. Treatment of Epstein-Barr-virus-positive post-transplantation lymphoproliferative disease with partly HLA-matched allogeneic cytotoxic T cells. The Lancet. 2002;360(9331):436-42.

14. Cho SG, Kim N, Sohn HJ, Lee SK, Oh ST, Lee HJ, et al. Long-term Outcome of Extranodal NK/T Cell Lymphoma Patients Treated With Postremission Therapy Using EBV LMP1 and LMP2a-specific CTLs. Molecular Therapy. 2015;23(8):1401–9.

15. Gottschalk S, Edwards OL, Sili U, Huls MH, Goltsova T, Davis AR, et al. Generating CTLs against the subdominant Epstein-Barr virus LMP1 antigen for the adoptive immunotherapy of EBV-associated malignancies. Blood. 2003;101(5):1905–12.

16. Ganesh K, Stadler ZK, Cercek A, Mendelsohn RB, Shia J, Segal NH, et al. Immunotherapy in colorectal cancer: rationale, challenges and potential. Nature Reviews Gastroenterology & Hepatology. 2019;16(6):361–75.

17. Coghill AE, Pfeiffer RM, Proietti C, Hsu W-L, Chien Y-C, Lekieffre L, et al. Identification of a novel, EBV-based antibody risk stratification signature for early detection of nasopharyngeal carcinoma in Taiwan. Clinical Cancer Research. 2018;24(6):1305–14.

18. Liu Z, Coghill AE, Pfeiffer RM, Proietti C, Hsu W-L, Chien Y-C, et al. Patterns of interindividual variability in the antibody repertoire targeting proteins across the Epstein-Barr virus proteome. The Journal of infectious diseases. 2018;217(12):1923–31.

19. Liu Z, Jarrett RF, Hjalgrim H, Proietti C, Chang ET, Smedby KE, et al. Evaluation of the antibody response to the EBV proteome in EBV-associated classical Hodgkin lymphoma. International Journal of Cancer. 2020;147(3):608–18.

20. Coghill AE, Proietti C, Liu Z, Krause L, Bethony J, Prokunina-Olsson L, et al. The association between the comprehensive Epstein–Barr virus serologic profile and endemic Burkitt lymphoma. Cancer Epidemiology and Prevention Biomarkers. 2020;29(1):57–62.

21. Liu Z, Sarathkumara YD, Chan JKC, Kwong Y-L, Lam TH, Ip DKM, et al. Characterization of the humoral immune response to the EBV proteome in extranodal NK/T-cell lymphoma. Scientific Reports. 2021;11(1):23664.

22. Proietti C, Zakrzewski M, Watkins TS, Berger B, Hasan S, Ratnatunga CN, et al. Mining, visualizing and comparing multidimensional biomolecular data using the Genomics Data Miner (GMine) Web-Server. Scientific Reports. 2016;6(1):1–15.

23. R Core Team R. R: A language and environment for statistical computing. 2013.

24. Biau G, Scornet E. A random forest guided tour. TEST. 2016;25(2):197–227.

25. Liaw A, Wiener M. Classification and regression by randomForest. R news. 2002;2(3):18–22.

26. Raskov H, Orhan A, Christensen JP, Gögenur I. Cytotoxic CD8+ T cells in cancer and cancer immunotherapy. British Journal of Cancer. 2021;124(2):359–67.

27. Huang J, Huang X, Huang J. CAR-T cell therapy for hematological malignancies: Limitations and optimization strategies. Frontiers in Immunology. 2022;13.

28. Anna M, Riccardo T, Riccardo D, Debora M, Elena M, Patrizia C, et al. The interplay between Epstein-Barr virus and the immune system: a rationale for adoptive cell therapy of EBV-related disorders. Haematologica. 2010;95(10):1769–77.

29. Hashmi AA, Hussain ZF, Hashmi KA, Zafar MI, Edhi MM, Faridi N, et al. Latent membrane protein 1 (LMP1) expression in Hodgkin lymphoma and its correlation with clinical and histologic parameters. World J Surg Oncol. 2017;15(1):89.

30. Salahuddin S, Fath EK, Biel N, Ray A, Moss CR, Patel A, et al. Epstein-Barr Virus Latent Membrane Protein-1 Induces the Expression of SUMO-1 and SUMO-2/3 in LMP1-positive Lymphomas and Cells. Scientific Reports. 2019;9(1):208.

31. Zeng M, Chen Y, Jia X, Liu Y. The Anti-Apoptotic Role of EBV-LMP1 in Lymphoma Cells. Cancer Manag Res. 2020;12:8801–11.

32. Pavlova S, Feederle R, Gärtner K, Fuchs W, Granzow H, Delecluse H-J. An Epstein-Barr Virus Mutant Produces Immunogenic Defective Particles Devoid of Viral DNA. Journal of Virology. 2013;87(4):2011–22.

33. Visalli RJ, Schwartz AM, Patel S, Visalli MA. Identification of the Epstein Barr Virus portal. Virology. 2019;529:152–9.

34. Chen LW, Hung CH, Wang SS, Yen JB, Liu AC, Hung YH, et al. Expression and regulation of the BKRF2, BKRF3 and BKRF4 genes of Epstein-Barr virus. Virus Res. 2018;256:76–89.

35. Tsurumi T, Kobayashi A, Tamai K, Daikoku T, Kurachi R, Nishiyama Y. Functional expression and characterization of the Epstein-Barr virus DNA polymerase catalytic subunit. Journal of virology. 1993;67(8):4651–8.

36. Okuno Y, Murata T, Sato Y, Muramatsu H, Ito Y, Watanabe T, et al. Defective Epstein-Barr virus in chronic active infection and haematological malignancy. Nat Microbiol. 2019;4(3):404–13.

37. Li J, Walsh A, Lam TT, Delecluse HJ, El-Guindy A. A single phosphoacceptor residue in BGLF3 is essential for transcription of Epstein-Barr virus late genes. PLoS Pathog. 2019;15(8):e1007980.

38. Liu X, Cohen JI. Epstein-Barr Virus (EBV) Tegument Protein BGLF2 Promotes EBV Reactivation through Activation of the p38 Mitogen-Activated Protein Kinase. Journal of Virology. 2016;90(2):1129–38.

39. Hung C-H, Chiu Y-F, Wang W-H, Chen L-W, Chang P-J, Huang T-Y, et al. Interaction Between BGLF2 and BBLF1 Is Required for the Efficient Production of Infectious Epstein– Barr Virus Particles. Frontiers in Microbiology. 2020;10.

40. Yap LF, Wong AKC, Paterson IC, Young LS. Functional Implications of Epstein-Barr Virus Lytic Genes in Carcinogenesis. Cancers. 2022;14(23):5780.

41. Tsurumi T, Kobayashi A, Tamai K, Yamada H, Daikoku T, Yamashita Y, et al. Epstein-Barr virus single-stranded DNA-binding protein: purification, characterization, and action on DNA synthesis by the viral DNA polymerase. Virology. 1996;222(2):352–64.

42. Xu M, Yao Y, Chen H, Zhang S, Cao SM, Zhang Z, et al. Genome sequencing analysis identifies Epstein-Barr virus subtypes associated with high risk of nasopharyngeal carcinoma. Nat Genet. 2019;51(7):1131–6.

43. Liao G, Huang J, Fixman ED, Hayward SD. The Epstein-Barr virus replication protein BBLF2/3 provides an origin-tethering function through interaction with the zinc finger DNA binding protein ZBRK1 and the KAP-1 corepressor. J Virol. 2005;79(1):245–56.

44. Van Esser JWJ, Niesters HGM, Thijsen SFT, Meijer E, Osterhaus ADME, Wolthers KC, et al. Molecular quantification of viral load in plasma allows for fast and accurate prediction of response to therapy of Epstein–Barr virus-associated lymphoproliferative disease after allogeneic stem cell transplantation. British Journal of Haematology. 2001;113(3):814–21.

45. Doubrovina E, Oflaz-Sozmen B, Prockop SE, Kernan NA, Abramson S, Teruya-Feldstein J, et al. Adoptive immunotherapy with unselected or EBV-specific T cells for biopsy-proven EBV+ lymphomas after allogeneic hematopoietic cell transplantation. Blood. 2012;119(11):2644–56.

46. Kim SS, Sumner WA, Miyauchi S, Cohen EEW, Califano JA, Sharabi AB. Role of B Cells in Responses to Checkpoint Blockade Immunotherapy and Overall Survival of Cancer Patients. Clinical Cancer Research. 2021;27(22):6075–82.

47. DeFalco J, Harbell M, Manning-Bog A, Baia G, Scholz A, Millare B, et al. Non-progressing cancer patients have persistent B cell responses expressing shared antibody paratopes that target public tumor antigens. Clinical Immunology. 2018;187:37–45.

48. Bruno TC, Ebner PJ, Moore BL, Squalls OG, Waugh KA, Eruslanov EB, et al. Antigen-Presenting Intratumoral B Cells Affect CD4+ TIL Phenotypes in Non–Small Cell Lung Cancer PatientsTIL-Bs Present Antigen to CD4 TILs in NSCLC. Cancer Immunology Research. 2017;5(10):898–907.

49. Kinker GS, Vitiello GAF, Ferreira WAS, Chaves AS, Cordeiro de Lima VC, Medina TDS. B Cell Orchestration of Anti-tumor Immune Responses: A Matter of Cell Localization and Communication. Frontiers in Cell and Developmental Biology. 2021;9:678127.

50. Shalapour S, Font-Burgada J, Di Caro G, Zhong Z, Sanchez-Lopez E, Dhar D, et al. Immunosuppressive plasma cells impede T-cell-dependent immunogenic chemotherapy. Nature. 2015;521(7550):94–8.

51. Kessel A, Haj T, Peri R, Snir A, Melamed D, Sabo E, et al. Human CD19+CD25high B regulatory cells suppress proliferation of CD4+ T cells and enhance Foxp3 and CTLA-4 expression in T-regulatory cells. Autoimmunity Reviews. 2012;11(9):670–7.

52. Khan AR, Hams E, Floudas A, Sparwasser T, Weaver CT, Fallon PG. PD-L1hi B cells are critical regulators of humoral immunity. Nature Communications. 2015;6(1):1–16.

53. Affara NI, Ruffell B, Medler TR, Gunderson AJ, Johansson M, Bornstein S, et al. B cells regulate macrophage phenotype and response to chemotherapy in squamous carcinomas. Cancer cell. 2014;25(6):809–21.

54. Shalapour S, Lin X-J, Bastian IN, Brain J, Burt AD, Aksenov AA, et al. Inflammation-induced IgA+ cells dismantle anti-liver cancer immunity. Nature. 2017;551(7680):340–5.

55. Ammirante M, Luo J-L, Grivennikov S, Nedospasov S, Karin M. B-cell-derived lymphotoxin promotes castration-resistant prostate cancer. Nature. 2010;464(7286):302–5.

56. Bruno TC. New predictors for immunotherapy responses sharpen our view of the tumour microenvironment. Nature. 2020;577(7791):474-6.

57. Tan GW, Visser L, Tan LP, van den Berg A, Diepstra A. The Microenvironment in Epstein-Barr Virus-Associated Malignancies. Pathogens. 2018;7(2).

58. Ellyard JI, Simson L, Parish CR. Th2-mediated anti-tumour immunity: friend or foe? Tissue Antigens. 2007;70(1):1–11.

59. Andreu-Sanz D, Kobold S. Role and Potential of Different T Helper Cell Subsets in Adoptive Cell Therapy. Cancers (Basel). 2023;15(6).

